# Evolution of reproductive traits have no apparent life-history associated cost in populations of *Drosophila melanogaster* selected for cold shock resistance

**DOI:** 10.1101/2020.04.22.047753

**Authors:** Karan Singh, Nagaraj Guru Prasad

**Affiliations:** Indian Institute of Science Education and Research Mohali, Knowledge City, Sector 81, SAS Nagar, PO Manauli, Punjab 140306, India, 2240124, 2240124

**Keywords:** Reproductive traits, life-history evolution, trade-off, life time fecundity, egg viability and mating number

## Abstract

In insect species like *Drosophila melanogaster*, the ability to evolve greater resistance or evolution of certain traits under specific environmental conditions leads to energy trade-offs with other important life-history traits. A number of studies from multiple fields have documented the life-history associated cost. However, no known studies have assessed the life-history associated cost with evolved reproductive traits and increase egg viability in cold shock selected population. To explore this, we used replicate populations of *D. melanogaster* that have evolved reproductive traits and egg viability in response to increased resistance to non lethal cold shock. To assess life-history cost; we measured longevity, life time fecundity, Larvae to adults development time, and larvae to adults survival. We found that there were no significant differences in longevity, life time fecundity, larvae to adults survival, and male body weight between the selected and control populations. However, selected populations have significantly longer pre adults developmental time compared to their control population. Females from the selected populations were bigger in size compared to the control populations. These findings suggest that there is no life-history cost associated with the evolution of greater resistance in the aspect of faster recovery of egg viability and reproductive traits post cold-shock. It quite possible the cost of the evolution of reproductive traits and egg viability in response to cold shock resistant is paid in terms of reduced resistance to other stresses

## INTRODUCTION

A number of ecological factors including temperature are known to vary across time and space and as a result, organisms experience different types of unfavorable environmental conditions during their lifespan. These environmental stresses can be major drivers of evolution of life-history of organisms in nature (Hoffman and Parsons, 1991, reviewed in Parsons, 2005).

Temperature is one of the fundamental ecological features of an organism’s environment. Organisms can respond to extreme temperatures in various ways, like changes in their behavior patterns and physiology or life-history traits (Hoffmann and Parsons, 1991, Patton and Krebs, 2001, Fasolo and Krebs, 2004). Resources used for coping with stress are unavailable for other functions under limited resource conditions, which can lead to trade-offs across important life-history traits such as somatic maintenance and reproduction (Stearns, 1992). For example, one important way in which organisms cope with immediate changes in temperature (heat-shock and cold-shock) is by expressing heat shock proteins (HSPs). Expression of these proteins is extremely costly and is known to affect reproduction (Krebs and Loeschcke, 1994). Thus, temperature shock can affect various important life-history traits (Huey and Berrigan, 2001, Hochachka and Somero, 2002, Sinclair et al., 2003, Angilletta, 2009). Deviation from ambient temperature (where absolute fitness of an organism is maximum) drastically affects various life-history and related traits of insects (Lee and Denlinger, 1991, Voituron et al., 2002, Hoffmann et al., 2003) such as fecundity, male fertility, lifespan (Denlinger and Yocum, 1998, Bubliy and Loeschcke, 2005, Rohmer et al, 2004, reviewed in Hance, 2007, Lieshout et al., 2013, Nguyen et al., 2013, Singh et al., 2016), reproduction (Singh et al., 2016, Singh and Prasad, 2016), mating ability (Singh et al., 2015), development time (Trotta et al., 2006, Austin and Moehring, 2013) and motility (Angilletta et al., 2002)

Several studies have investigated the evolution of life-history traits in response to thermal variation. *D. melanogaster* being widely distributed offers a great model to study the evolution of life-history traits in response to temperature variation across latitudes and altitudes. In general, a number of traits vary progressively across populations inhabiting various latitudes. This pattern of results suggests that life-history evolution in populations of *Drosophila* is primarily being driven by environmental differences and that the populations are adapting to local environment, most probably, including temperature, which is an important component of the environment. Latitudinal clines have been found in a number of life-history traits such as development time, survivorship, larval competitive ability, fecundity and body size (Stanley and Parsons, 1981, Bouletreau-Merle et al., 1982, James and Partridge, 1995, 1998, reviewed in Hoffmann et al., 2003, Hangartner et al., 2015).

Some experimental evolution studies have investigated the evolution of life-history traits in response to selection for cold stress tolerance (Tucic, 1979, Chen and Walker, 1993, Watson and Hoffmann, 1996, Anderson et al., 2005, Bubliy and Loeschcke 2005, MacMillan et al., 2009). However, such studies are a few (Overgaard et al., 2010). Anderson et al. (2005) found increased female fecundity and decreased male longevity in populations of *D. melanogaster* selected for rapid chill-coma recovery. MacMillan et al. (2009) also documented reduced longevity in females (but not in males) in populations selected for increased resistance to freeze shock. However, Bubliy and Loeschcke (2005) did not notice any difference in longevity and development time in populations of *D. melanogaster* selected for increased cold tolerance. Thus, the correlated evolution of life-history traits in response to cold stress has been fairly variable.

In this study, our aim in probing the life-history costs, if any, of increased resistance to cold stress in terms of increased reproductive traits and egg viability (Singh et al., 2015, Singh et al. 2016, Singh and Prasad 2016) in populations of *D. melanogaster* selected for cold shock resistance. To investigate the underlying life-history cost to increased resistance to cold stress, we assayed various life-history (longevity and life time fecundity) and related traits such as development time, and body weight in the populations of *D. melanogaster* selected for increased resistance to cold stress. These experiments were performed over 24-33 generations of selection.

## MATERIALS AND METHODS

### Experimental populations

Details of the maintenance and derivation of the selected (FSB; Cold shock Selected population derived from BRB population) and their control (FCB; Cold shock Control derived from BRB population) populations has been explained previously (Singh et al. 2015). Briefly, after 35 generation of the laboratory adaption of BRB 1-5, one FSB population and one FCB population were established from each of the BRB populations, for example, FSB 1 and their corresponding control FCB 1 derived from the BRB 1, similarly FSB 2 and their control FCB 2 established from the BRB2 and so on. Hence, we had five replicate populations for the selected population and five replicate of control populations. Populations carrying the same numerical subscript have originated from same base line population (BRB) and are more close to each other than any other populations. For instance, FSB 1 and FCB 1 are more close (due to the origin from the same ancestral population) to each other than FSB 2 or FCB 2 or any other population. Hence, in our statistical data analysis FSB 1 and FCB 1 are included in block 1, similarly FSB 2 and FCB 2 are included in block 2 and so on. FSB and FCB populations are large outbred populations maintained under standard laboratory environment (25°C temperature, 50–60% relative humidity, 12 hour light: 12 hour dark cycle, on a 13-day discrete generation cycle). On day 12 post egg collection, flies (flies are roughly 2-3 days old as adults and mated) are moved into empty, clean, dry glass vials (30 mm diameter × 90 mm length). After that flies belonging to the FSB populations are subjected to - 5°C temperature in ice-salt-water slurry for one hour. FCB populations, on the other hand, are held at 25°C for one hour. Subsequently, all populations are quickly moved into a separate Plexiglass cage (25 cm length × 20 cm width × 15 cm height) having a fresh food plate. After 24 hour a fresh food plate is given to flies in order to collect eggs to initiate next generation. For each population, 20 vials are collected at density of 70 eggs per vial containing ∼ 6 ml of a fresh food.

### Standardization

To account the non-genetic parental effects (Rose 1984), flies from the selected populations and from their controls were reared for one generation in common rearing environment. This method is referred to as standardization and these flies are known as standardized flies. A detail of the standardization of the protocol has been described earlier in Singh et al. (2015). Shortly, to control eggs density, for each selected and their control populations, 20 vials were established at density of 70 eggs per vials in ∼6 ml food, reared at standard laboratory conditions (12 hours light:12 hours dark). On day 12 after egg collection, (roughly 2-3 days old as adult flies) ∼ 1200-1400 flies of each population were transferred separately in a Plexiglass cage and provided a fresh food plate. These flies were further used for experiment egg collection.

### Cold shock treatment for experiments

Detailed account of the cold shock protocol has been described in our previous study (Singh et al., 2015). In short, on day 12 post egg collection, (by this time flies were roughly 2-3 days old as adult and mated flies) 25 pairs of males and females were moved to clean, dry glass vials under mild carbon dioxide anesthesia. The cotton plug was inserted deep into the vial such that the flies were allowed to stay in a confined space in vial (1/3 of the vial). The flies were kept in an incubator to recover from carbon dioxide anesthesia for half an hour. The vials containing flies were then kept for one hour in ice-salt-water slurry maintained at -5°C. Post cold shock, flies were quickly shifted to Plexiglass cages (14 cm length × 16 cm width × 13 cm height. The cage was provided with a food Plate and was kept under standard laboratory conditions (Singh et al., 2015). The control treatment flies were handled similar way, except that the vials containing flies, were kept in a water bath that maintained at 25°C for one hour.

## Experimental details

### Experiment: 1.1: Longevity assay

The longevity assay was performed after 24 generations of selection. Eggs were collected from standardized flies at a controlled egg density of 70 eggs/vial provisioned with ∼ 6 ml of fresh banana-yeast-jaggery food (hereafter referred to as “food”). Twenty four such type of vials were set up for each of the FSB (1-5) and FCB (1-5) populations. On day 12 after egg collection, flies were sorted (25 mating pairs per vial) under mild carbon dioxide anesthesia. After sorting, flies were divided into two sets: (a) set first for cold-shock treatment (both male and female flies were exposed to cold shock for one hour) and (b) set second for no-shock treatment (neither males and nor females were exposed to cold-shock).

#### (a)Cold-shock

For each population, flies contained in 12 vials (each vial contains 25 mating pairs of male and female) were imposed cold-shock (−5°C for one hour) as mentioned in the cold shock protocol. Quickly, after the cold shock, 12 vials were randomly divided into 3 sets referred to as a “replicate”. Each set having 4 vials of flies (100 mating pairs each) were moved into a Plexiglass cage and given a fresh food plate. Hence, each population (FSB 1-5 and FCB 1-5) had 3 replicates.

#### (b)No-shock

For each population, flies contained in 12 vials (each vial contain 25 mating pairs of male and female) were subjected to no-shock treatment (25°C for one hour). Post treatment, 12 vials were quickly randomly divided into three sets that were known as replicate. Each set having 4 vials containing total of 100 mating pairs of male and female flies were moved into Plexiglas cages and given a fresh food plate. Hence, each population (FSB 1-5 and FCB 1-5) had 3 replicates.

We established three replicate cages per selection **×** block **×** treatment combination (Except block 1 of the FCB population which had 2 replicates for both-shock treatment, due to accidental death of one of the replicates during the assay). Food plate was changed 48 hours internal and dead flies were aspirated out and computed. Sex of the dead flies was determined under microscope on the basis of sex combs. Mortality was recorded until the last fly died. Using the mortality data, for each cage, we measured mean longevity of males and females from the selection regime (FSB and FCB), treatment and block. For the analysis of mean longevity, cage means were used as the unit of analysis.

### Experiment 1.2: Life time fecundity assay

Fecundity assay was performed along with longevity assay, using the same set of flies. We measured fecundity at every sixth day along with longevity. In order to measure fecundity, fresh food plate was placed in the Plexiglas cage for 6 hours for oviposition. After that, total number of eggs on each plate was counted under the microscope. Subsequently, fecundity per female - number of eggs divided by total number of live females at that time point-was calculated. Average fecundity of the eleven time points was calculated for three replicates for each of the FSB 1-5 and FCB 1-5 populations, and treatments. We also computed median and maximum longevity for each cage populations. The aging rate data was analyzed using the Gompertz model.

### Experiment 2: Development time (first instar larva to eclosion)

Development time was assayed after 33 generations of selection. Followed by one generation of common rearing environment or standard laboratory condition (no selection was imposed on FSB and FCB population), 12 vials each were set up for FSB 1-5 and FCB 1-5 populations at a density of 70 eggs per vial. On day 12 after egg collection, vials containing flies were randomly divided into two sets for - (a) cold-shock (b) no-shock treatment. For both ‘cold-shock’ and ‘no-shock’ treatments, flies were transferred into empty glass vials at density of ∼70 flies and the cotton plug was pushed deep up to the bottom one-third of the vial. After that, the flies were subjected to cold shock or no shock treatments, following the protocol as mentioned above. Immediately after cold-shock treatment, flies (200 males and 200 females) were transferred to Plexiglass cage and provided with a fresh food plate. Twenty four hours post cold-shock, fresh - food plates were given to each cage for 1 hour to lay stored eggs. After that another set of fresh plates were given for four hours. The second set of plates containing eggs were then incubated at standard laboratory conditions for 18 hours to allow eggs to hatch and first instars larvae to emerge. The larvae were collected (using a moist brush) into vials with 6 ml of a fresh food. For each population and treatment combination, 10 replicate vials were set up (each containing 30 larvae in 6 ml of food). The vials were incubated at standard laboratory conditions. The positions of the vials were randomized and moved daily within the incubator. Once pupae formed, each vial was manually scanned every 2 hours. Freshly eclosed flies were transferred into empty glass vials, sexed and counted. The flies were then flash frozen using liquid nitrogen and then transferred to -80°C for storage used to assess dry body weight.. Mean larva to eclosion development time was computed for each vial and this vial mean time was considered as the unit of analysis.

### Experiment 3: Measurement of dry body weight of male and female flies

In order to measure the dry body weight, we used the same flies from the development time assay (mentioned above). Freshly eclosed flies were flash frozen using liquid nitrogen and stored at -80°C until dry body weight measurement. Five flies of a given sex were grouped together, dried in a hot air oven at 65°C for 48 hours and weighed. For each population, treatment and sex combination, ten such sets were weighed. Thus, a total of 50 males and 50 females per population and treatment were used for body weight measurement. Body weight of each group of five flies was considered as the unit of analysis.

### Experiment 4: Larvae to adults survival

To investigate larvae to adults survival, we monitored total number of flies eclosed from the cultured larval vial at density of 30 larvae/vial. We calculated percentage of larvae to adult survival using the equation in given a bracket (percentage of larvae to adults survival = (number of eclosed flies in a vial/total number of larvae cultured in a vial)*100).

### Statistical analysis

Mean longevity, development time, dry body weight of males and females, and larvae to adults survival were analyzed using a three-factor mixed model analysis of variance (ANOVA) treating selection regime (FSB vs. FCB), treatment (cold-shock vs. no-shock) as a fixed factors crossed with a random block (1-5). The sexes were analyzed separately.

Fecundity per female was analyzed using a three-factor mixed model ANOVA treating selection regime (FSB vs. FCB) and treatment (cold-shock vs. no-shock) as fixed factors crossed with block as a random factor. All the analyses were done at α=0.05 level of significance using Statistica (for Windows, version 10, Statsoft). Multiple comparisons were carried out employing Tukey’s HSD.

### Rates of aging

Age dependent and age independent rate of aging was measured using the method used by Mueller et al. (1995) and Jafariet al. (2007). Raw survivorship data were used to calculate ‘proportion survival’ values with subsequent calculation of running average of the proportion survival data, *r*_*x*_.

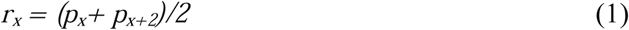

Where, *p*_*x*_ is the proportion of individuals surviving at a given age *x*. Since mortality was monitored every alternate day, *x* and *x+2* are two successive age intervals noticed. The hazard rate that is the probability of death per unit time, μ_*x*_at age x was computed employing the following equation:

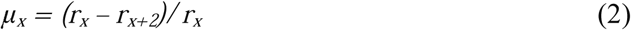

According to the Gompertz equation, the mortality rate at age *x* is given by,

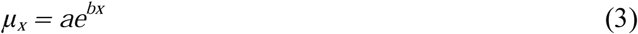

Where, *a* and *b* represent age-independent and age-dependent rate of aging respectively. Log-hazard rate was regressed against age intervals; the intercept and the least square slope gave the estimates of *Gompertz a* and *Gompertz b* respectively. The derived parameters were analyzed using three factor mixed model ANOVA with selection regime (FSB vs. FCB), treatment (cold shock vs. no shock) as fixed factor crossed with random blocks (1-5).

## RESULTS

### Experiment 1.1: Longevity assay for male and female

Male and female longevity was assessed in terms of mean, median and maximum longevity. Analyses revealed that the results were similar regardless of the measure used. After 24 generations of selection, there was no significant effect of selection, treatment or selection × treatment interaction on male or female mean longevity (Table 1a, b Figure 1a, b, c, and d). Interestingly, the absence of any significant effect of treatment indicated that flies subjected to cold-shock treatment as well as the flies that were not subjected to cold-shock, which proved that cold shock had no direct effect on mean longevity.

**Table 1a.** Effect of cold-shock on the mean longevity of males (Experiment 1.1).Summary of results from a three-factor mixed model ANOVA on the male mean longevity using selection (FCB and FSB) and treatment (cold-shock and no-shock) as fixed factors crossed with random block (1-5).

| Effect | SS | MS Num | DF Num | DF Den | F ratio | p |
| --- | --- | --- | --- | --- | --- | --- |
| Selection (Sel) | 33.439 | 33.439 | 1.000 | 4.006 | 1.348 | 0.310 |
| Treatment (Trt) | 14.549 | 14.549 | 1.000 | 4.008 | 0.752 | 0.435 |
| Block (Blk) | 478.234 | 119.559 | 4.000 | 6.177 | 3.024 | 0.107 |
| Sel × Trt | 9.169 | 9.169 | 1.000 | 4.032 | 1.972 | 0.232 |
| Sel × Blk | 99.283 | 24.821 | 4.000 | 4.000 | 5.350 | 0.067 |
| Trt × Blk | 77.398 | 19.349 | 4.000 | 4.000 | 4.170 | 0.098 |
| Sel × Trt × Blk | 18.559 | 4.640 | 4.000 | 39.000 | 0.422 | 0.792 |

**Table 1b.** Effect of cold-shock on the mean longevity of females (Experiment 1.1).Summary of results from a three-factor mixed model ANOVA on the female mean longevity using selection (FCB and FSB) and treatment (cold-shock and no-shock) as fixed factors crossed with random block (1-5). *p*-values in bold are statistically significant.

| Effect | SS | MS Num | DF Num | DF Den | *F* ratio | *p* |
| --- | --- | --- | --- | --- | --- | --- |
| Selection (Sel) | 56.044 | 56.044 | 1.000 | 4.007 | 2.756 | 0.172 |
| Treatment (Trt) | 1.581 | 1.581 | 1.000 | 4.004 | 0.047 | 0.839 |
| Block (Blk) | 91.967 | 22.992 | 4.000 | 7.012 | 0.443 | 0.775 |
| Sel × Trt | 0.160 | 0.160 | 1.000 | 4.071 | 0.084 | 0.786 |
| Sel × Blk | 81.399 | 20.350 | 4.000 | 4.000 | 10.751 | **0.020** |
| Trt × Blk | 133.984 | 33.496 | 4.000 | 4.000 | 17.697 | **0.008** |
| Sel × Trt × Blk | 7.571 | 1.893 | 4.000 | 39.000 | 0.192 | 0.941 |

**Table 1c.** Effect of cold shock on the age independent and age dependent mortality rates of males (Experiment 1.1). Summary of results from a three-factor mixed model ANOVA on (a) age independent and (b) age dependent mortality rate among males using Selection (FCB and FSB) and Treatment (cold-shock and no-shock) as fixed factors crossed with random block (1-5). *p*-values in bold are statistically significant.

| Trait | Effect | SS | MS Num | DF Num | DF Den | *F* ratio | *p* |
| --- | --- | --- | --- | --- | --- | --- | --- |
| (a) Male | Selection (Sel) | 0.479 | 0.479 | 1.000 | 4.021 | 2.920 | 0.162 |
| age | Treatment (Trt) | 9.979 | 9.979 | 1.000 | 4.021 | 61.398 | **0.001** |
| independent | Block (Blk) | 8.791 | 2.198 | 4.000 | 0.008 | 126.988 | 0.960 |
| Mortality | Sel × Trt | 0.127 | 0.127 | 1.000 | 4.011 | 0.410 | 0.557 |
|  | Sel × Blk | 0.656 | 0.164 | 4.000 | 4.000 | 0.531 | 0.723 |
|  | Trt × Blk | 0.649 | 0.162 | 4.000 | 4.000 | 0.525 | 0.726 |
|  | Sel × Trt × Blk | 1.236 | 0.309 | 4.000 | 39.00 | 1.220 | 0.318 |
| (b) Male | Selection (Sel) | 3.9×10<sup>-5</sup> | 3.9×10<sup>-5</sup> | 1.000 | 4.017 | 0.340 | 0.591 |
| age | Treatment (Trt) | 7.3×10<sup>-3</sup> | 7.3×10<sup>-3</sup> | 1.000 | 4.030 | 112.215 | **<0.001** |
| dependent | Block (Blk) | 1.4×10<sup>-3</sup> | 3.5×10<sup>-4</sup> | 4.000 | 9×10<sup>-6</sup> | 3470.59 | 1.000 |
| Mortality | Sel × Trt | 1.8×10<sup>-9</sup> | 1.8×10<sup>-9</sup> | 1.000 | 4.011 | 1×10<sup>-5</sup> | 0.998 |
|  | Sel × Blk | 4.6×10<sup>-4</sup> | 1.1×10<sup>-4</sup> | 4.000 | 4.000 | 0.638 | 0.663 |
|  | Trt × Blk | 2.6×10<sup>-4</sup> | 6.4×10<sup>-5</sup> | 4.000 | 4.000 | 0.362 | 0.825 |
|  | Sel × Trt × Blk | 7.1×10<sup>-4</sup> | 1.8×10<sup>-4</sup> | 4.000 | 39.00 | 1.251 | 0.305 |

**Table 1d.**
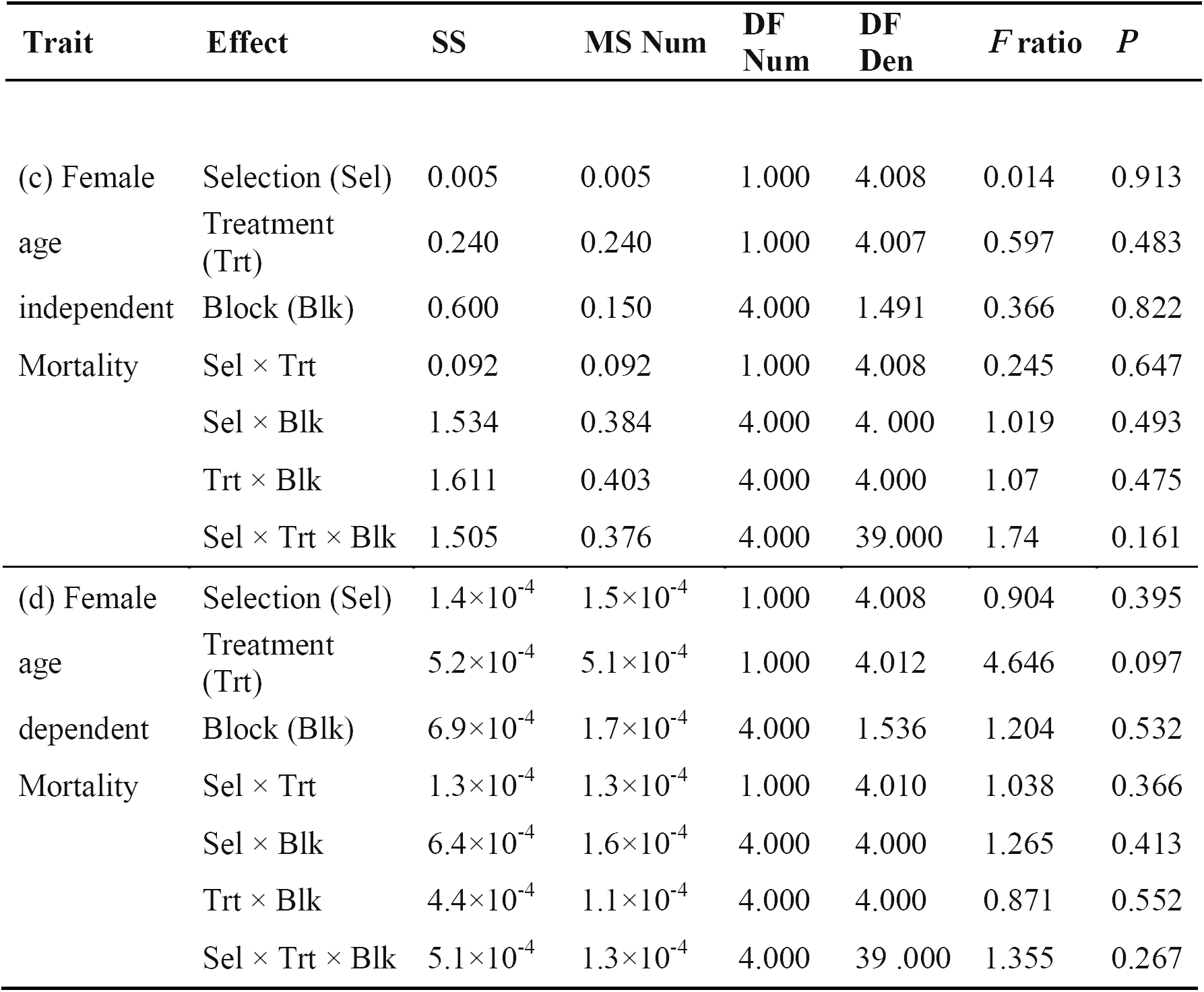
Effect of cold-shock on the age independent and age dependent mortality rates of females (Experiment 1.1).Summary of results from a three-way mixed model ANOVA on (c) age independent and (d) age dependent mortality rate among females using selection (FCB and FSB) and treatment (cold-shock and no-shock) as fixed factors crossed with random block (1-5).

**Figure 1a:**
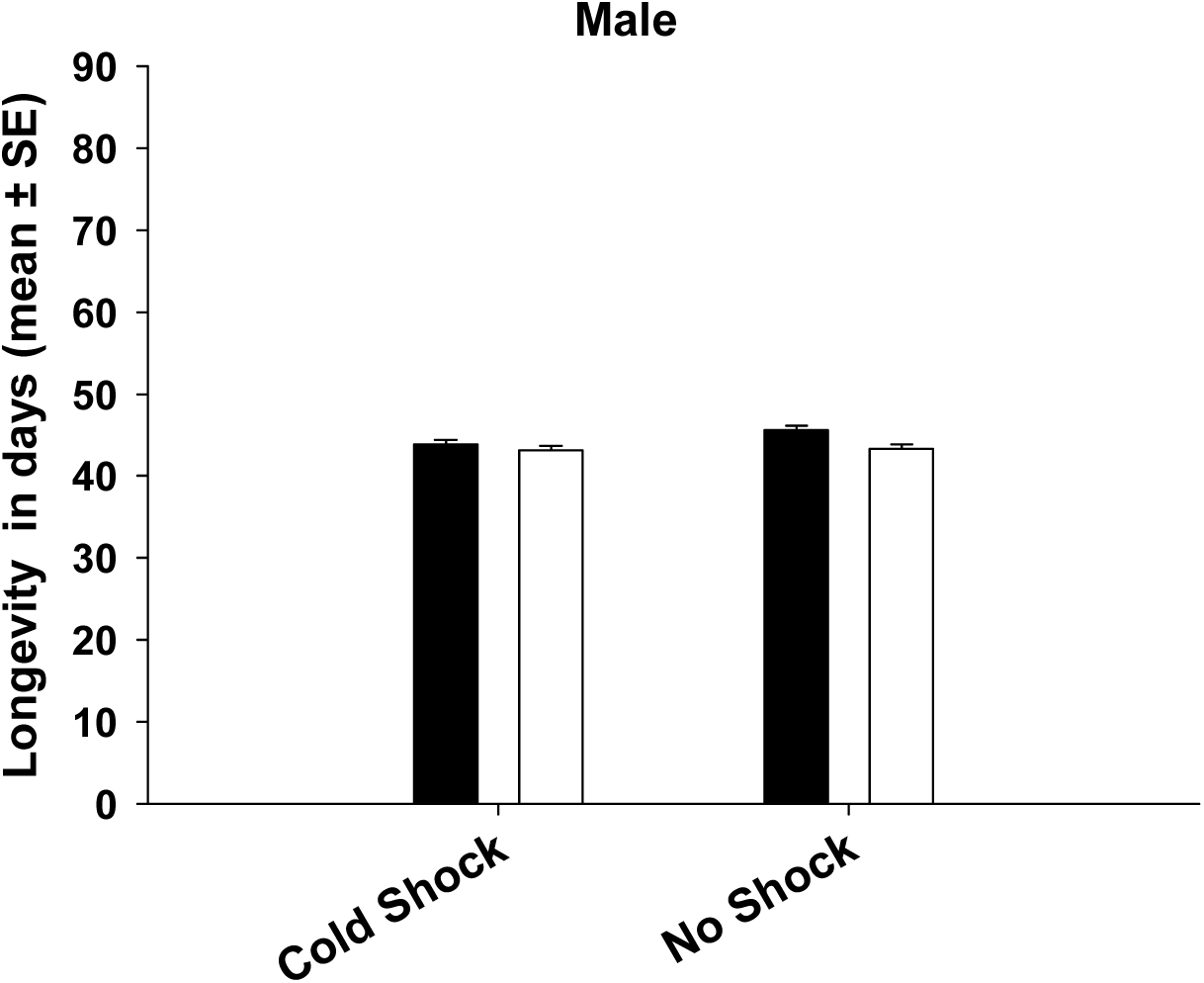
Mean longevity of the FSB and FCB males after being subjected to cold-shock or no-shock treatment (Experiment 1.1). Selection, treatment or selection × treatment interaction did not have significant effect on male mean longevity. Open bars represent the FSB and closed bars represent the FCB populations.

**Figure 1b:**
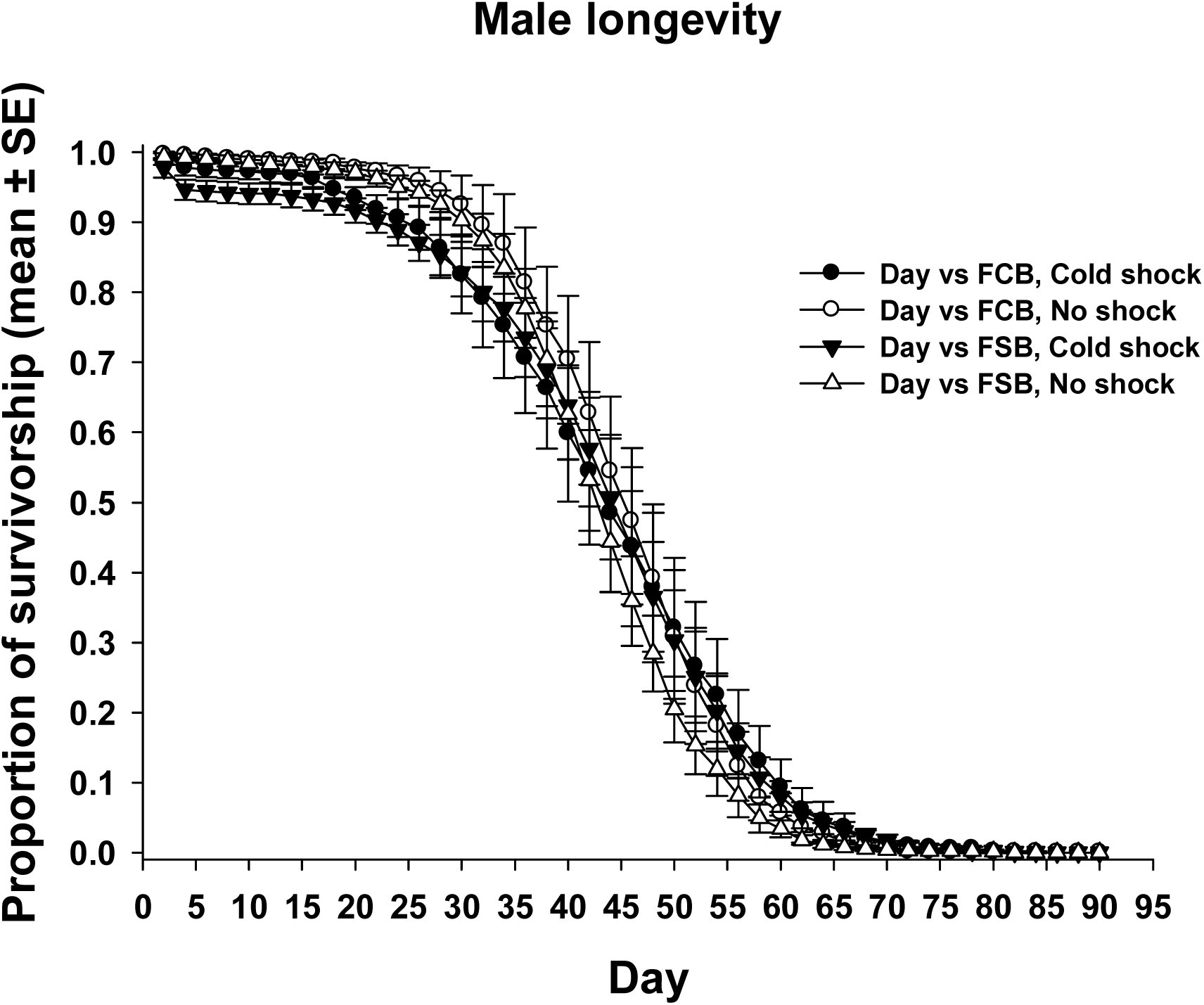
Male survivorship across ages (Experiment 1.1). Longevity was assayed after adult flies were subjected to cold-shock or no-shock treatment. There was no difference in the mean, median and maximum longevity of the FSB and FCB males. There was no significant difference in the *Gompertz* parameters between the FSB and FCB populations.

**Figure 1c:**
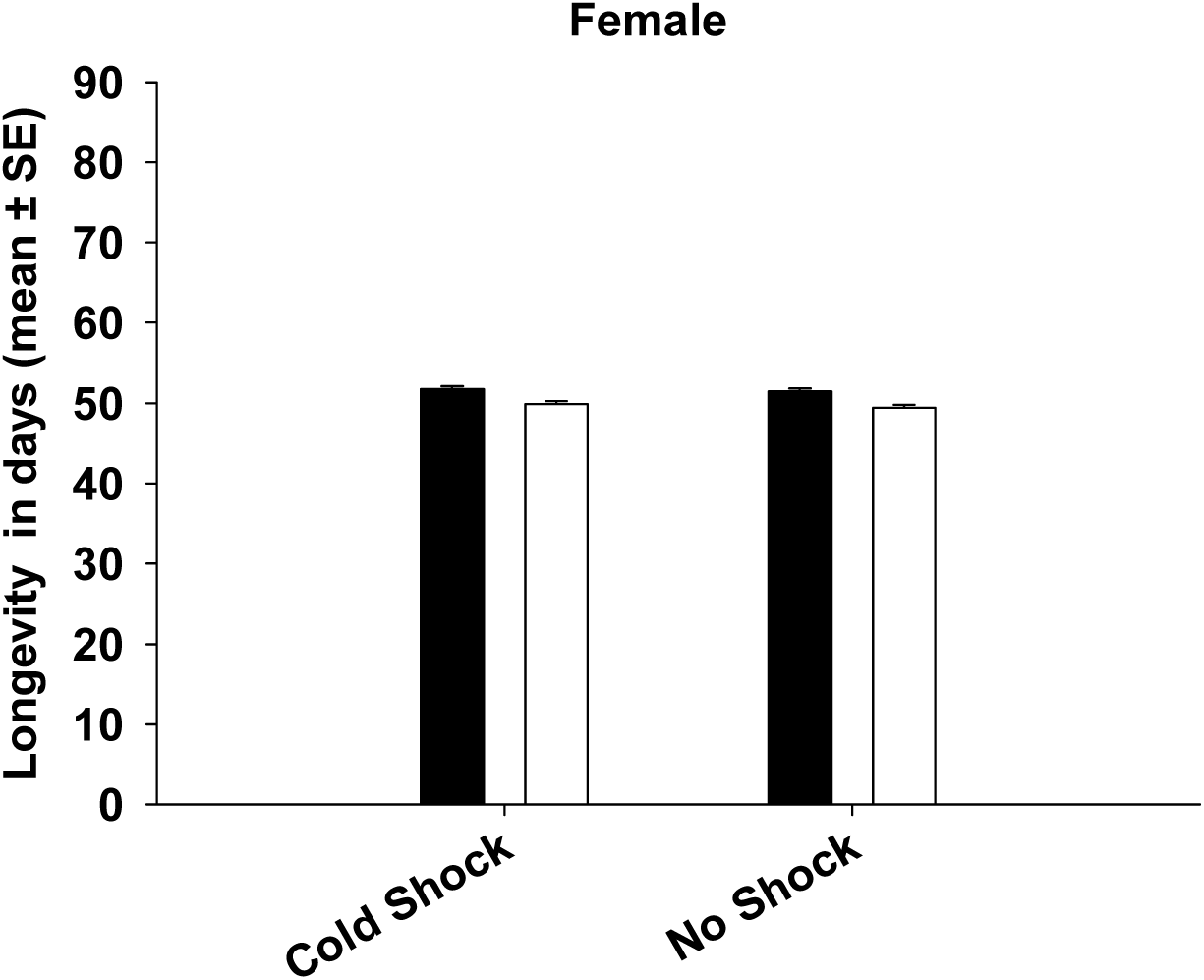
Mean longevity of the FSB and FCB females after being exposed to cold shock or no shock treatment (Experiment 1.1). Selection, treatment or selection × treatment interaction did not have significant effect on female mean longevity. Open bars represent the FSB and closed bars represent the FCB populations.

**Figure 1d:**
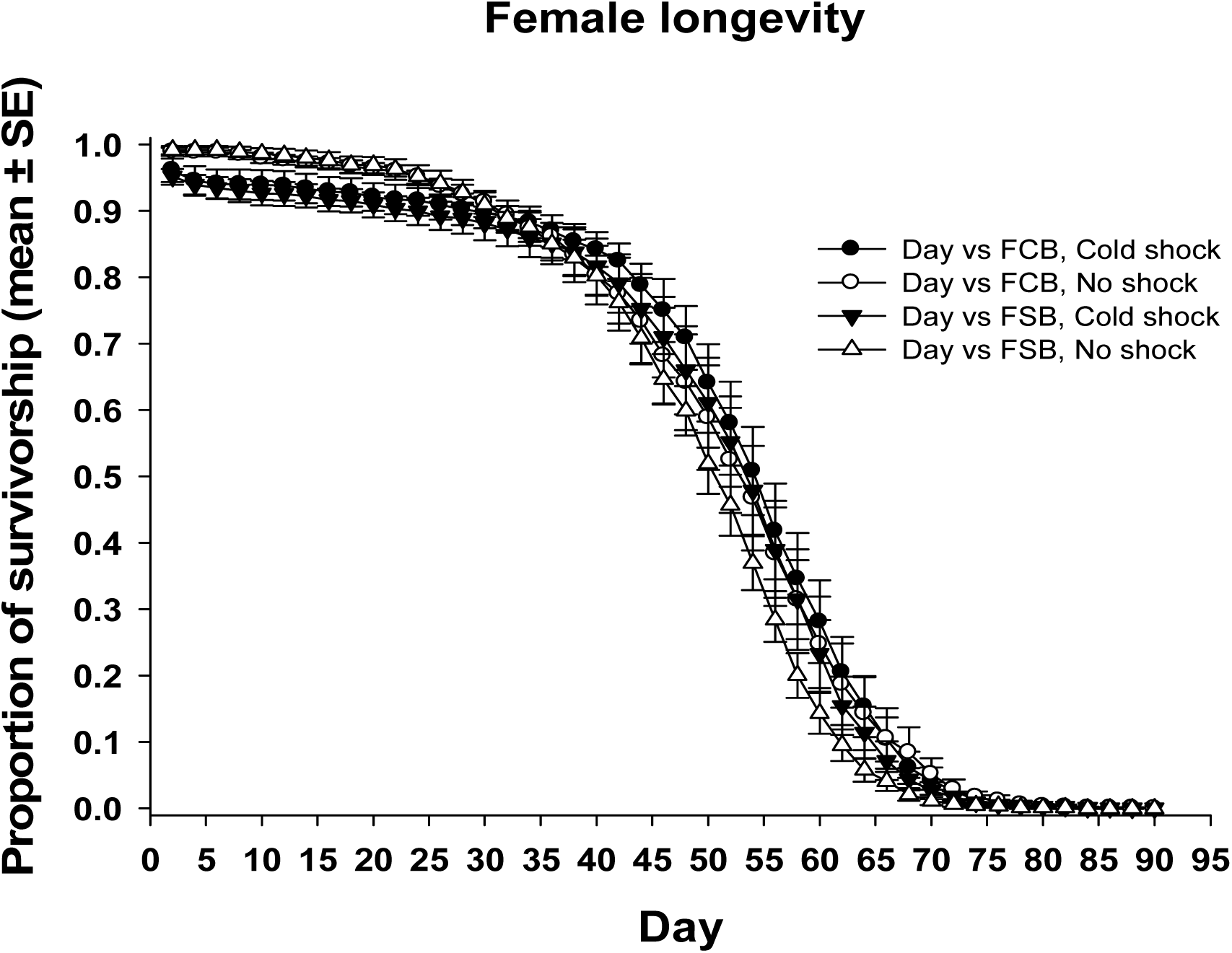
Female survivorship across ages (Experiment 1.1). Longevity was assayed after adult flies were subjected to cold-shock or no-shock treatment. There was no difference in the mean, median and maximum longevity of the FSB and FCB females. There was no significant difference in the *Gompertz* parameters between FSB and FCB populations.

We found a significant effect of treatment on the *Gompertz a* (age independent mortality rate) and b (age dependent mortality rate) parameters among males. The FSB and FCB males subjected to cold-shock showed significantly higher age independent mortality but a significantly lower age dependent mortality compared to the males not subjected to cold-shock (Table 1c). The net effect of these two factors was that the average (and median) lifespan of the males subjected to cold-shock and those not subjected to cold-shock was not different. There was no effect of selection or a selection × treatment interaction on the Gompertz parameters. Among the females, none of the factors affected the Gompertz parameters (Table 1d). Thus, we found no evidence for any significant change in mean longevity or rates of aging as a correlated response to selection for increased resistance to cold-shock.

### Experiment 1.2: Life time fecundity

The mean number of eggs laid per female in each of the FSB and FCB populations and treatments were computed by averaging across the 11 time points of fecundity measurement and used it as the unit of analysis. We did not observe significant effects of selection, treatment or selection × treatment interaction on female fecundity (Table 2, Figure 2a, and 2b). Just like longevity, the absence of any significant effect of treatment on fecundity revealed that cold-shock treatment had no direct effect on lifetime fecundity.

**Table 2.**
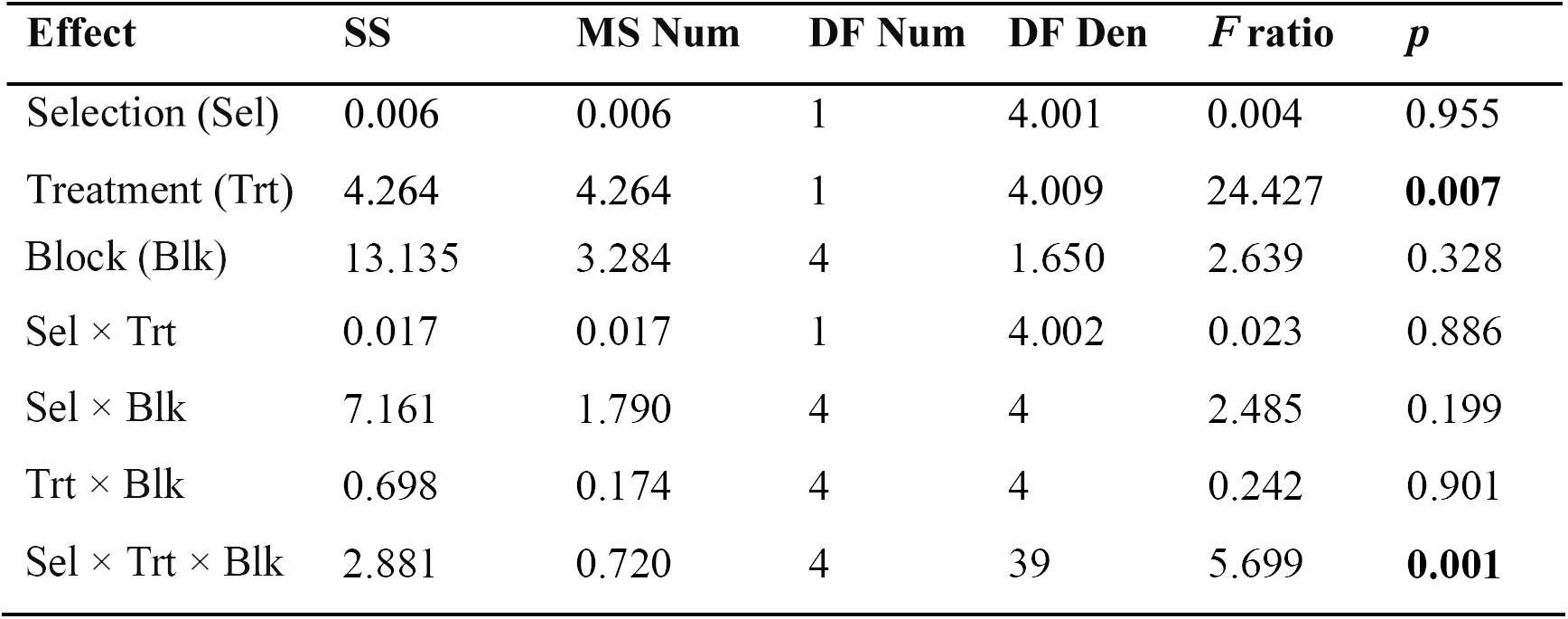
Effect of cold shock on life time fecundity (Experiment 1.2).Summary of results from a three-factor mixed model ANOVA on the life time fecundity using selection (FCB and FSB) and treatment (cold-shock and no-shock) as fixed factors crossed with random block (1-5).

**Figure 2a:**
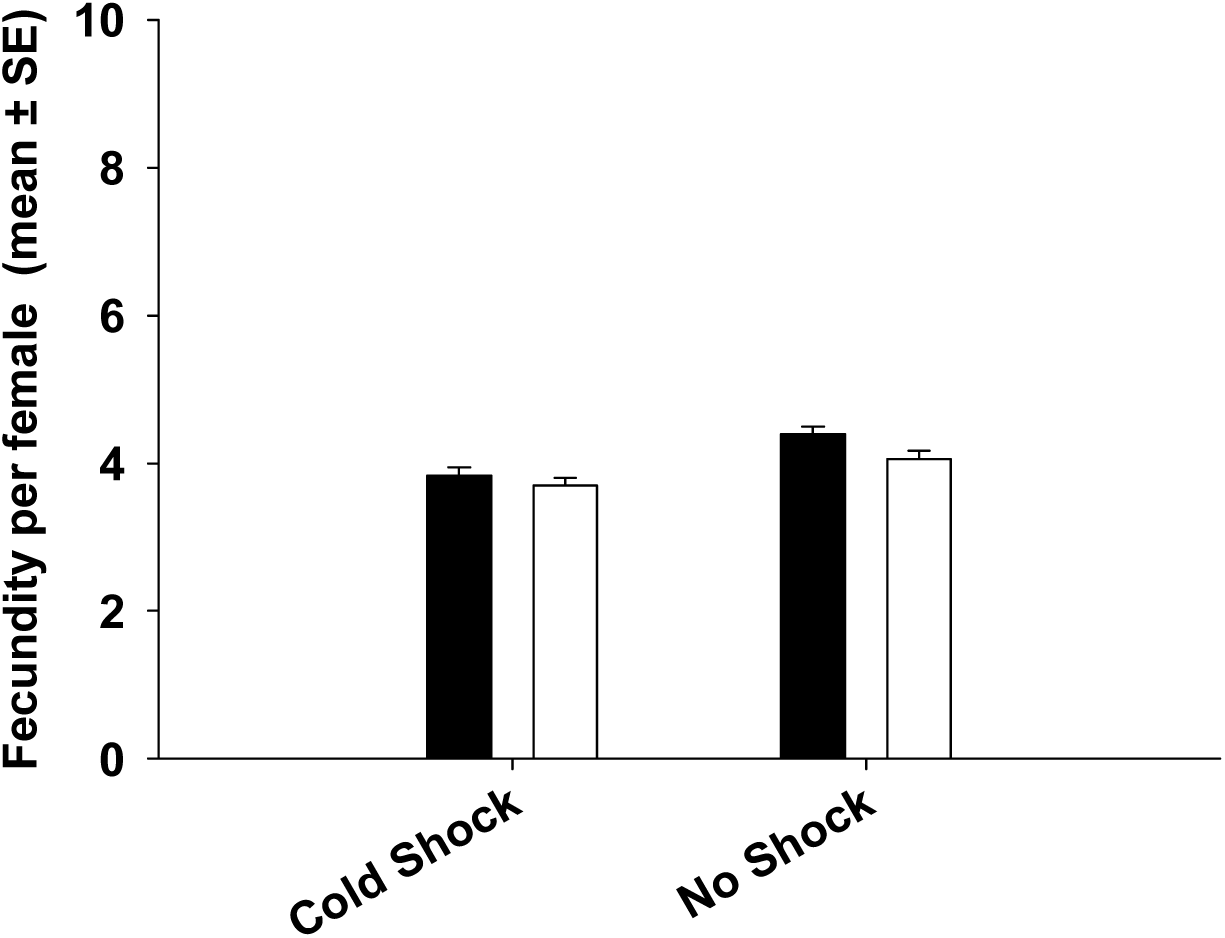
Mean life time fecundity per female (Experiment 1.2). Fecundity was measured at eleven time points once in every 6 days and mean of eleven time points for fecundity was computed. Selection, treatment or selection × treatment interaction did not have significant effect on fecundity. Open bars represent the FSB populations and closed bars represent the FCB populations.

**Figure 2b:**
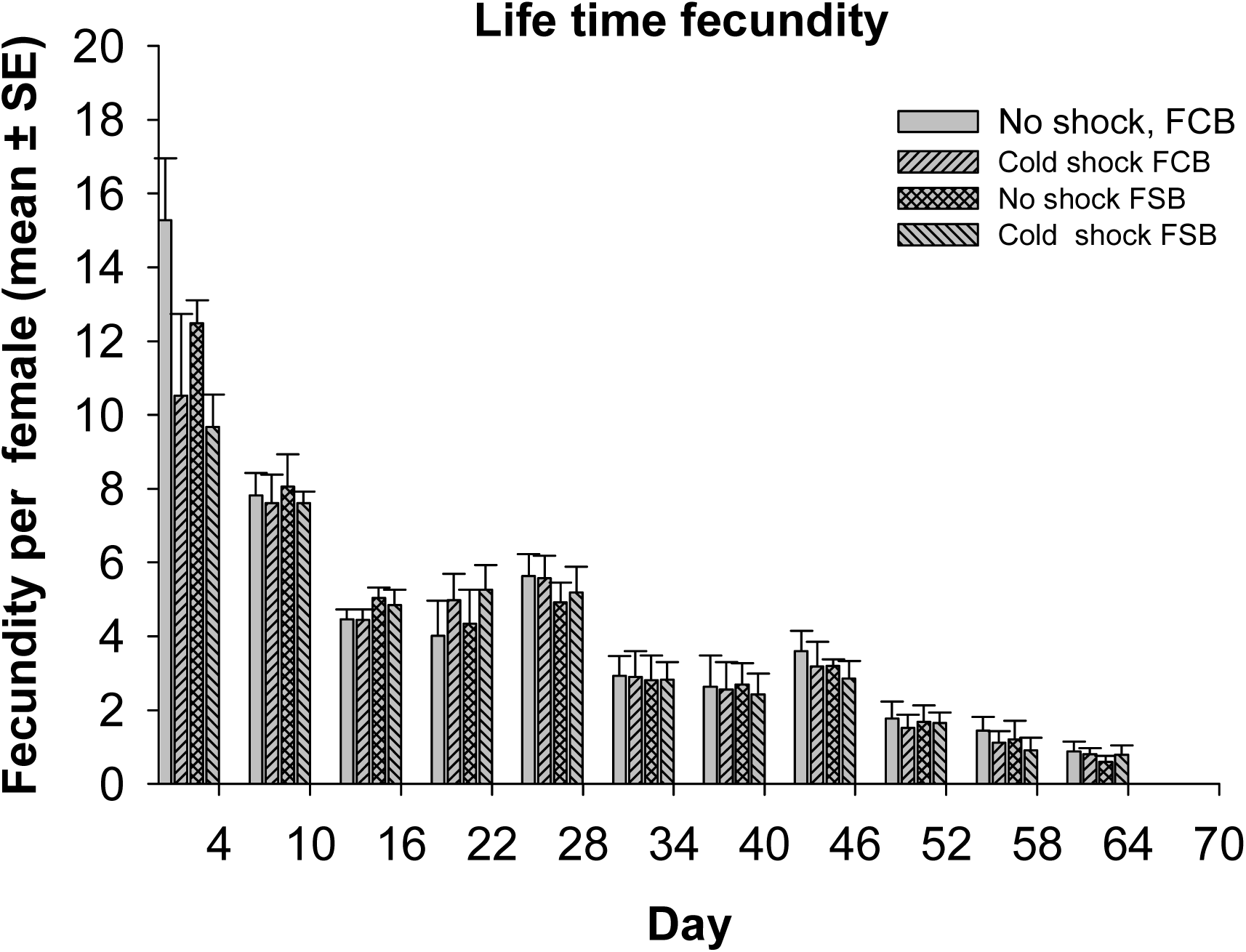
Life time fecundity per female (Experiment 1.2). Fecundity was measured at eleven time points once in every 6 days. Mean fecundity per female for each population and treatment was computed for eleven time points. Results indicate that fecundity reduces with age. However, none of the other effects were significant.

### Experiment 2: Development time (first instar larva to adult eclosion)

Unlike longevity and fecundity, selection did affect mean development time. Mean development time of males showed a significant effect of selection. (Table 3a, Figure 3a). Starting as first instar larvae, FSB male took about 2-4 hours more to emerge as adults compared to FCB males (Figure 3a). Female mean development time analysis showed that there was significant effect of selection (Table 3b, Figure3b). However, none of the other effects were significant. Just like the males, FSB females also took∼3-6 hours more to emerge as adults compared to FCB females (Figure3b). Again, the cold shock experienced by the parents had no effect on offspring development time (no significant treatment effect).

**Table 3a.** Effect of cold-shock on parents on the male developmental time (larvae to adult eclosion) (Experiment 2).Summary of results from a three-factor mixed model ANOVA on the mean larva to adult development time of males using selection (FCB and FSB) and treatment (cold-shock and no-shock) as fixed factors crossed with random block (1-5). *p*-values in bold are statistically significant.

| Effect | SS | MS Num | DF Num | DF Den | *F* ratio | *p* |
| --- | --- | --- | --- | --- | --- | --- |
| Selection (Sel) | 758.240 | 758.240 | 1.000 | 4.000 | 17.449 | **0.014** |
| Treatment (Trt) | 214.335 | 214.335 | 1.000 | 4.000 | 0.601 | 0.482 |
| Block (Blk) | 141.786 | 35.446 | 4.000 | 3.993 | 0.098 | 0.977 |
| Sel × Trt | 54.776 | 54.776 | 1.000 | 4.000 | 1.404 | 0.302 |
| Sel × Blk | 173.816 | 43.454 | 4.000 | 4.000 | 1.114 | 0.460 |
| Trt × Blk | 1427.513 | 356.878 | 4.000 | 4.000 | 9.150 | **0.027** |
| Sel × Trt × Blk | 156.012 | 39.003 | 4.000 | 180.000 | 1.111 | 0.353 |

**Table 3b.** Effect of cold-shock on parent on the female developmental time (larvae to adult eclosion) (Experiment 2).Summary of results from a three-factor mixed model ANOVA on mean larva to adult development time of females using selection (FCB and FSB) and treatment (cold-shock and no-shock) as fixed factors crossed with random block (1-5). *p*-values in bold are statistically significant.

| Effect | SS | MS Num | DF Num | DF Den | *F* ratio | *p* |
| --- | --- | --- | --- | --- | --- | --- |
| Selection (Sel) | 907.888 | 907.888 | 1.000 | 4.000 | 8.374 | **0.044** |
| Treatment (Trt) | 206.835 | 206.835 | 1.000 | 4.000 | 0.453 | 0.538 |
| Block (Blk) | 950.196 | 237.549 | 4.000 | 1.019 | 0.855 | 0.658 |
| Sel × Trt | 204.729 | 204.729 | 1.000 | 4.000 | 0.712 | 0.446 |
| Sel × Blk | 433.683 | 108.421 | 4.000 | 4.000 | 0.377 | 0.816 |
| Trt × Blk | 1828.026 | 457.007 | 4.000 | 4.000 | 1.590 | 0.332 |
| Sel × Trt × Blk | 1149.930 | 287.483 | 4.000 | 180.000 | 1.840 | 0.123 |

**Figure 3a:**
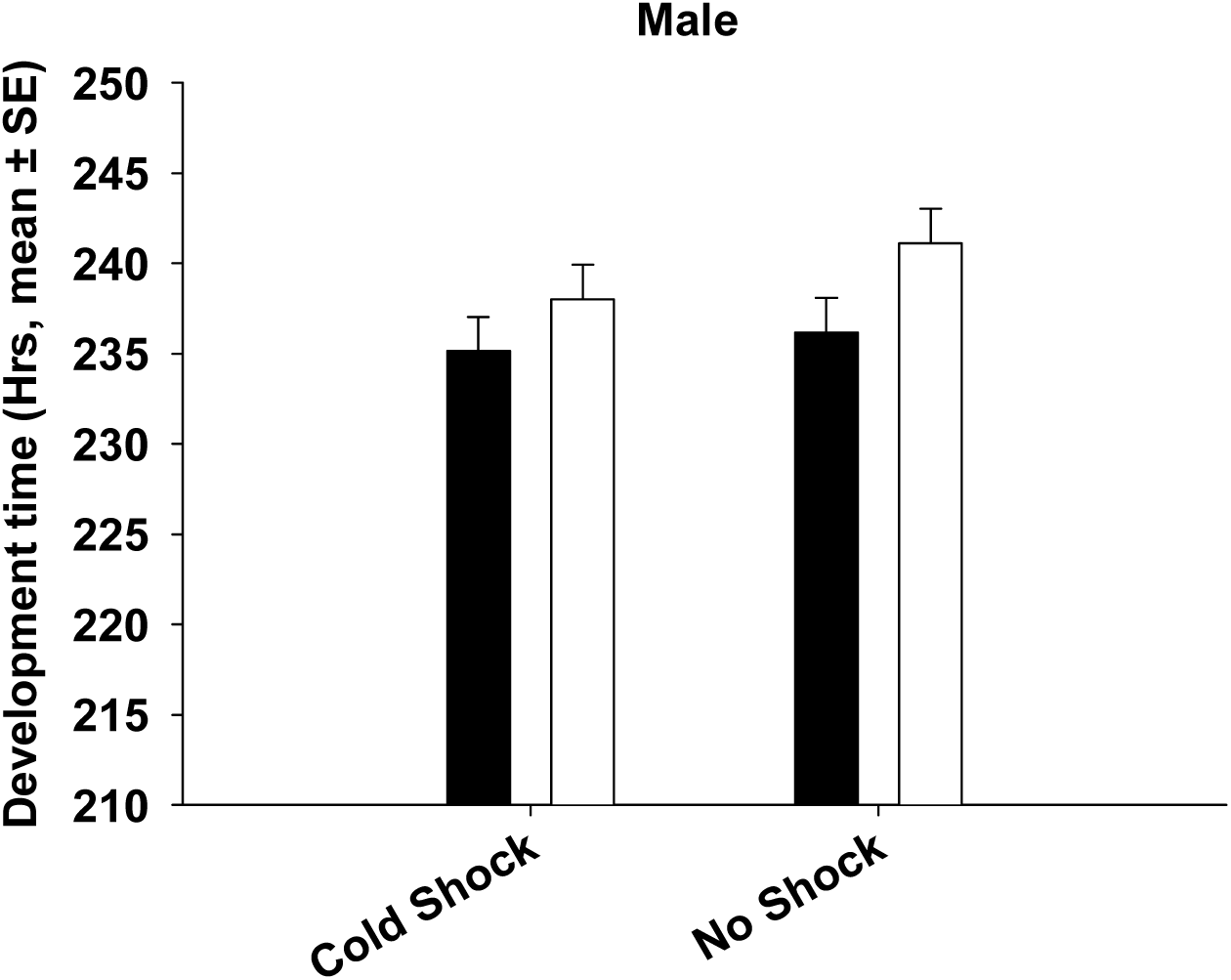
Mean development time (larva to adult) of the FSB and FCB males when their parents were subjected cold-shock or no-shock treatments (Experiment 2). We found a significant effect of selection regime with the FSB males developing 3-4 hours slower than FCB males. Treatment had no significant effect. Open bars represent the FSB populations and closed bars represent the FCB populations.

**Figure 3b:**
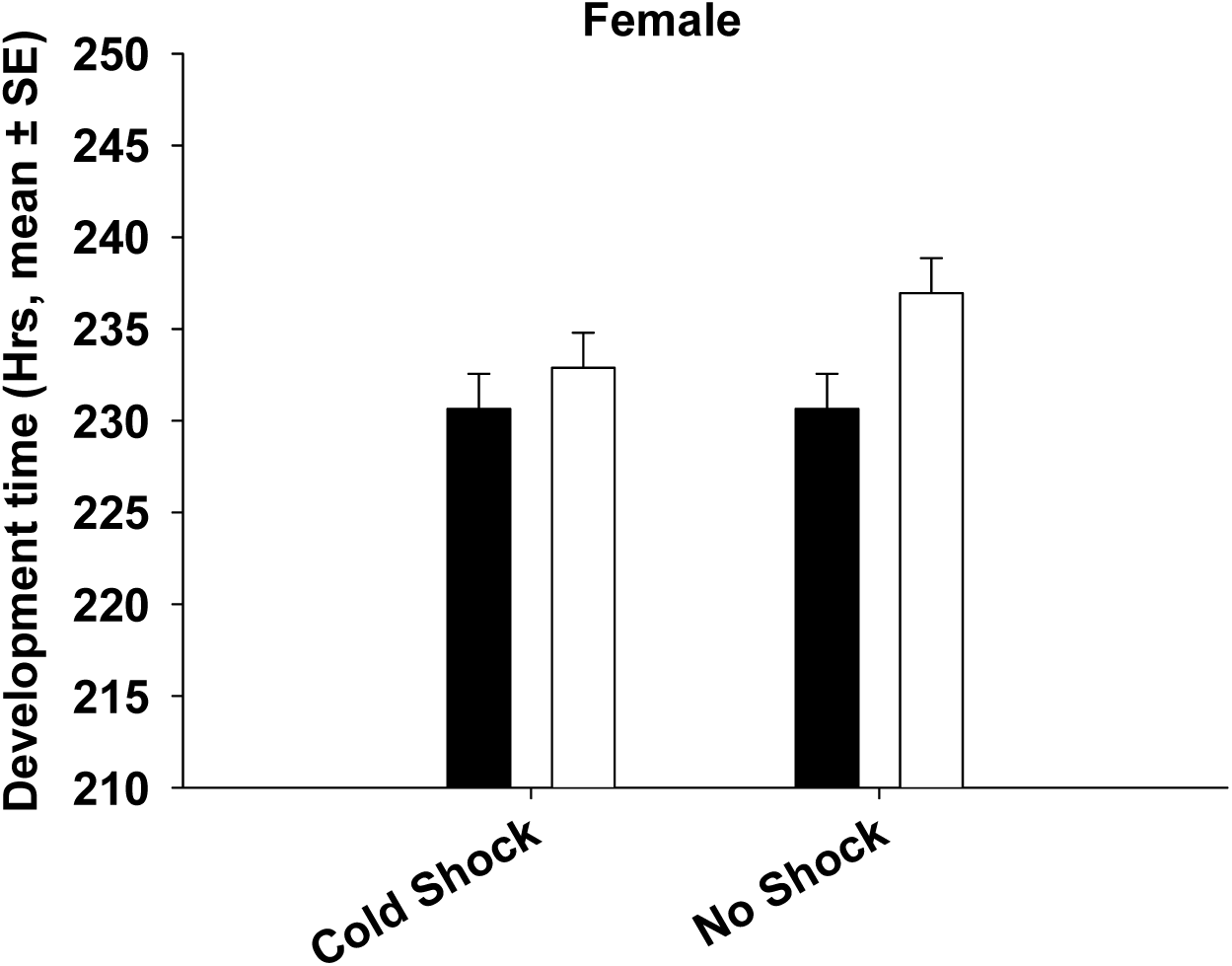
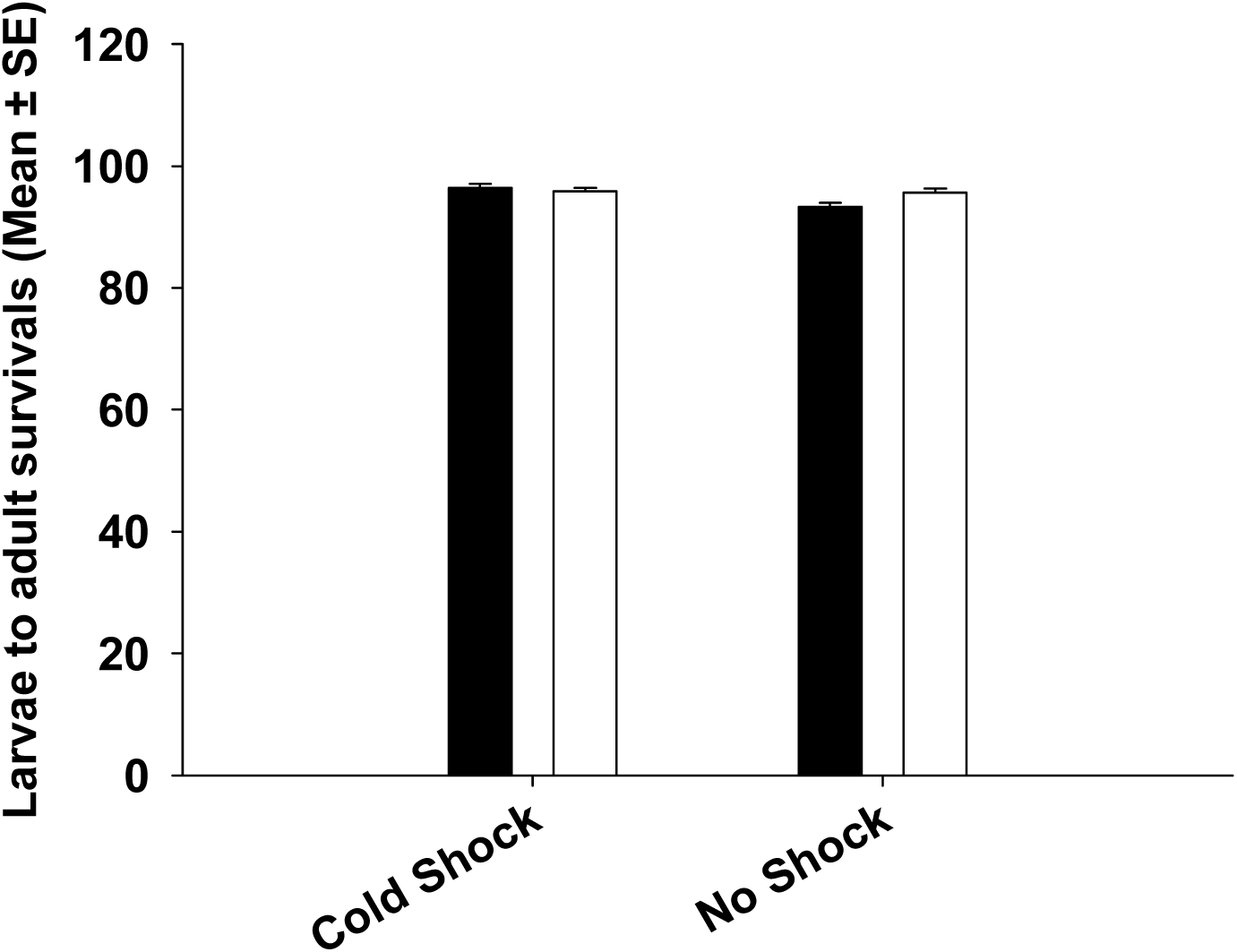
Mean development time (larva to adult) of the FSB and FCB females when their parents were subjected cold-shock or no-shock treatments (Experiment 2). We found a significant effect of Selection regime with FSB females developing 3-4 hours slower than FCB females. Treatment had no significant effect. Open bars represent the FSB populations and closed bars represent the FCB populations.

### Experiment 3: Dry body weight

Male mean dry body weight analysis revealed that there was no significant effect of selection, treatment or selection × treatment interaction (Table 4a, Figure 4a). In case of female dry body weight, found significant main effect of selection (Table 4b). However, there was no significant effect of treatment or selection × treatment interaction (Table 4b). Mean body weight of FSB females was about ∼0.01 mg higher than that of FCB females (Figure 4b).

**Table 4a.** Effect of cold-shock on the dry body weight of male (Experiment 3).Summary of results from a three-factor mixed model ANOVA on the mean dry body weight of males using selection (FCB and FSB) and treatment (cold-shock and no-shock) as fixed factors crossed with random block (1-5). *p*-values in bold are statistically significant.

| Effect | SS | MS Num | DF Num | DF Den | *F* ratio | *p* |
| --- | --- | --- | --- | --- | --- | --- |
| Selection (Sel) | 8.2×10<sup>-5</sup> | 8.2×10<sup>-5</sup> | 1.000 | 4.000 | 0.178 | 0.694 |
| Treatment (Trt) | 7×10<sup>-4</sup> | 7×10<sup>-4</sup> | 1.000 | 4.000 | 0.701 | 0.450 |
| Block (Blk) | 6×10<sup>-3</sup> | 1.5×10<sup>-3</sup> | 4.000 | 6.553 | 1.050 | 0.450 |
| Sel × Trt | 1.2×10<sup>-5</sup> | 1.2×10<sup>-5</sup> | 1.000 | 4.000 | 0.240 | 0.650 |
| Sel × Blk | 1.8×10<sup>-3</sup> | 4.6×10<sup>-4</sup> | 4.000 | 4.000 | 9.550 | **0.025** |
| Trt × Blk | 4×10<sup>-3</sup> | 1×10<sup>-3</sup> | 4.000 | 4.000 | 20.974 | **0.006** |
| Sel × Trt × Blk | 2×10<sup>-4</sup> | 4.8×10<sup>-5</sup> | 4.000 | 180.000 | 0.145 | 0.965 |

**Table 4b.** Effect of cold shock on the dry body weight of female (Experiment 3).Summary of results from a three-factor mixed model ANOVA on the mean dry body weight of females using selection (FCB and FSB) and treatment (cold-shock and no-shock) as fixed factors crossed with random Blocks (1-5). *p*-values in bold are statistically significant.

| Effect | SS | MS Num | DF Num | DF Den | *F* ratio | *p* |
| --- | --- | --- | --- | --- | --- | --- |
| Selection (Sel) | 6.7×10<sup>-3</sup> | 6.7×10<sup>-3</sup> | 1.000 | 4.000 | 32.942 | **0.005** |
| Treatment (Trt) | $3 \times 10^{-4}$ | $3 \times 10^{-4}$ | 1.000 | 4.000 | 0.287 | 0.621 |
| Block (Blk) | $1.4 \times 10^{-2}$ | $3.4 \times 10^{-3}$ | 4.000 | 3.756 | 3.620 | 0.128 |
| Sel × Trt | $2 \times 10^{-4}$ | $2 \times 10^{-4}$ | 1.000 | 4.000 | 1.199 | 0.335 |
| Sel × Blk | $8 \times 10^{-4}$ | $2 \times 10^{-4}$ | 4.000 | 4.000 | 1.059 | 0.479 |
| Trt × Blk | $3.7 \times 10^{-3}$ | $9 \times 10^{-4}$ | 4.000 | 4.000 | 4.828 | 0.078 |
| Sel × Trt × Blk | $8 \times 10^{-4}$ | $2 \times 10^{-5}$ | 4.000 | 180.000 | 0.491 | 0.743 |

**Figure 4a:**
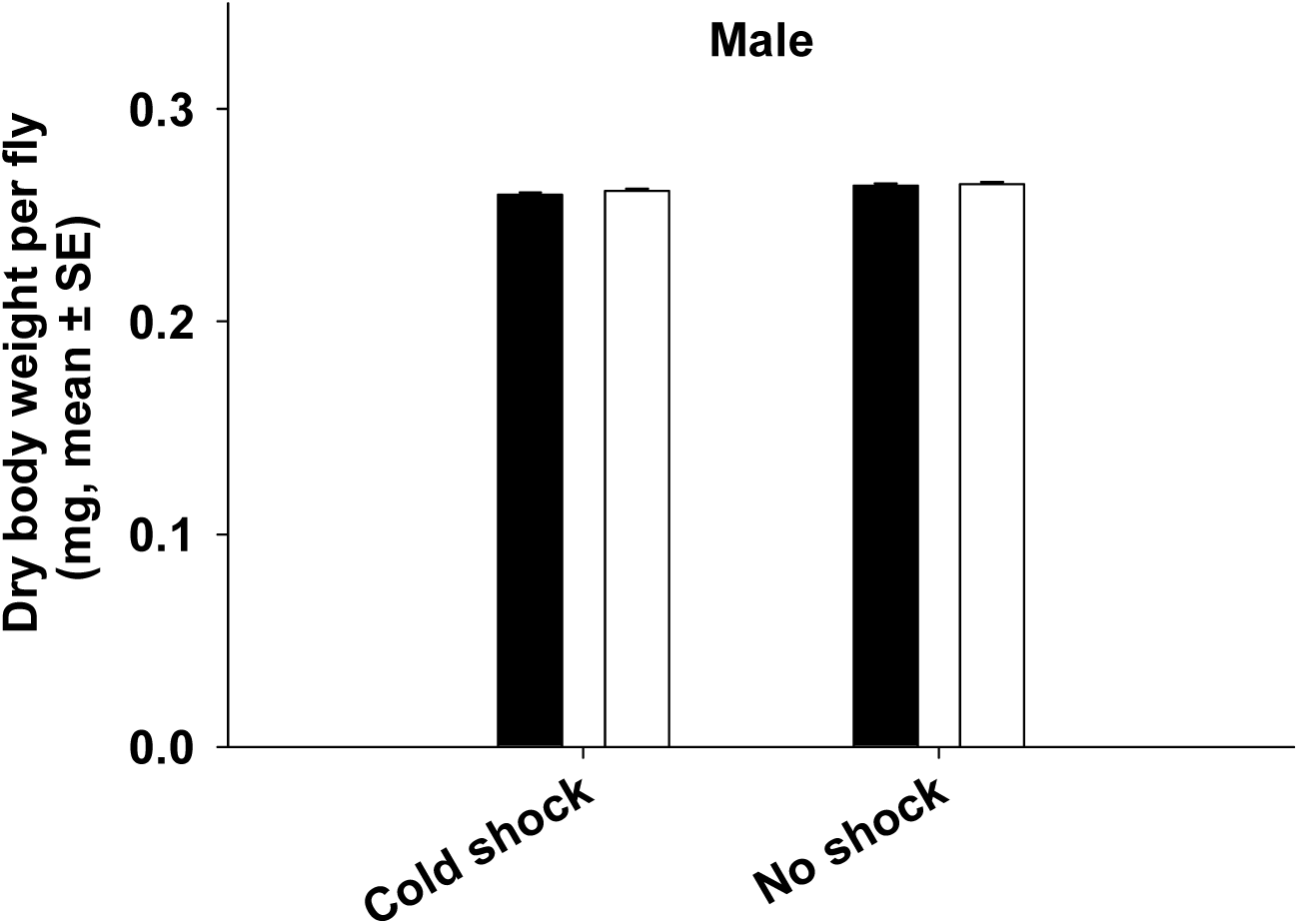
Dry weight at eclosion of males from the FSB and FCB populations (Experiment 3). Selection, treatment or selection × treatment interaction did not have significant effect on mean dry body weight. Open bars represent the FSB and closed bar represent the FCB populations.

**Figure 4b:**
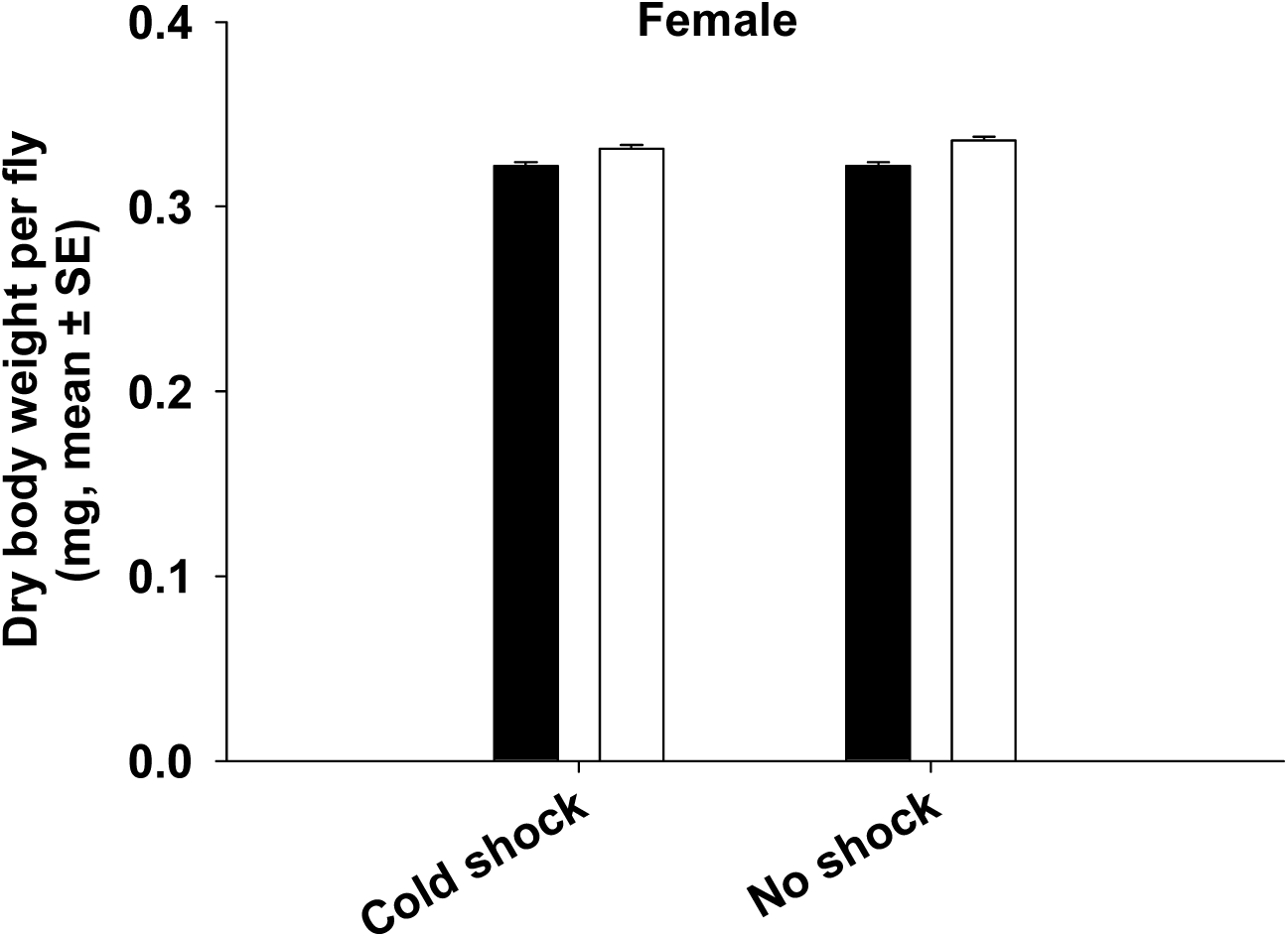
Dry weight at eclosion of females from the FSB and FCB populations. Selection had significant effect on mean dry body weight (Experiment 3). However, treatment or selection × treatment did not have significant effects on mean dry body weight. Open bars represent the FSB and closed bar represent the FCB populations.

### Experiment 4: larvae to adults survival

Mean larvae to adult survivals analysis shown that there was no significant effect of selection or selection × treatment interaction (Table 5, Figure 5).

**Table 5.**
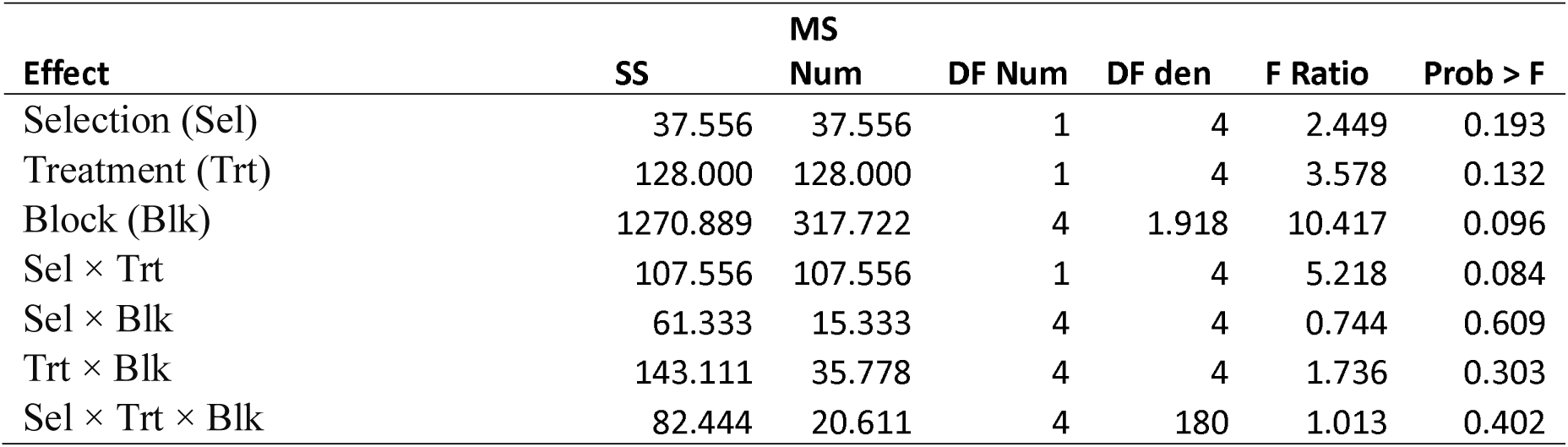
Effect of cold shock on the larvae to adults survivals (Experiment 4).Summary of results from a three-factor mixed model ANOVA on the mean larvae to adults survivals considering selection (FCB and FSB) and treatment (cold-shock and no-shock) as fixed factors crossed with random Blocks (1-5). *p*-values in bold are statistically significant.

**Figure 5:**
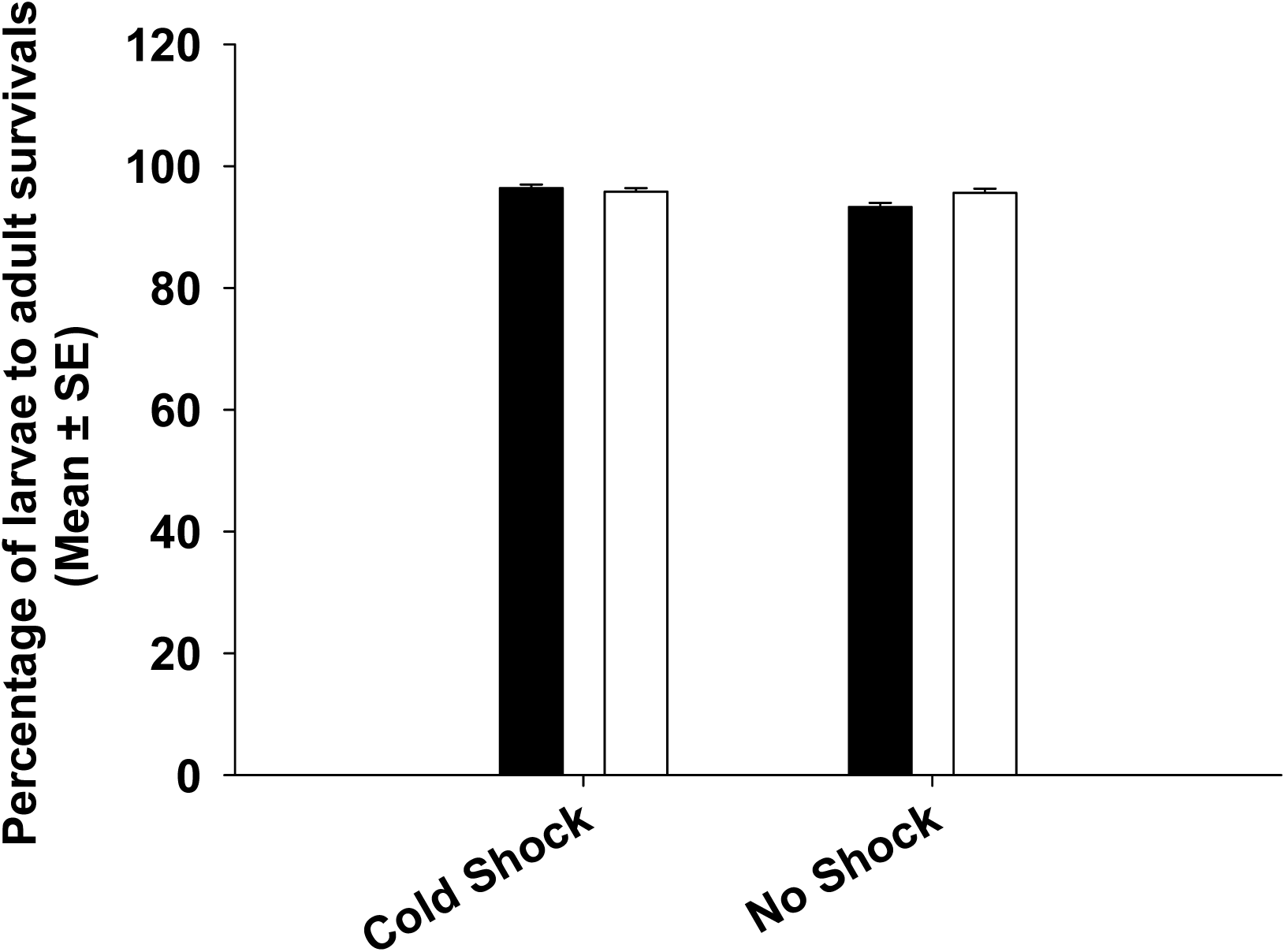
Larvae to adults survival from the FSB and FCB populations (Experiment 4). Selection, treatment or selection × treatment did not have any significant effects on mean larvae to adult survivals. Open bars represent the FSB and closed bar represent the FCB populations

## DISCUSSION

In this study, we assessed mean longevity, rates of aging, developmental time and dry body weight in the FSB and FCB populations with and without cold shock. Neither longevity nor fecundity was different between the FSB and FCB populations. However, we found that males and females from the FSB populations took significantly more time to develop (from first instar larvae to adult) relative to the FCB populations. Females from the FSB populations were heavier than females from the FCB populations. However, there was no difference in male body size between the FSB and FCB populations. Taken together, our finding suggests there is no evidence for a trade-off between the ability to resist cold stress and important life-history traits.

The correlation between cold-shock resistance and longevity is variable across studies. MacMillan et al. (2009), using a selection protocol very similar to the present study found that females of the cold-shock selected populations had decreased longevity compared to females of the control populations whereas no such difference was visible in the males. According to Anderson et al. (2005) populations selected for faster chill-coma recovery had reduced lifespan compared to controls. In the contrary Norry and Loeschcke (2002) observed that cold adapted populations lived longer at 14°C and shorter at 25°C compared to control populations. Bubliy and Loeschcke (2005) found no change in female longevity between populations selected for cold resistance and their controls. In populations directly selected for increased lifespan, increased cold resistance evolved as a correlated response in adults and pupae of *D. melanogaster* (Luckinbill, 1998). In contrast to all these studies, we found that selection for resistance to cold-shock had no effect on lifespan or rates of aging. There were several possible differences including the base population used for selection, the definition of ‘cold stress’, the assay protocols, *etc*. between these studies that preclude a direct comparison of results. More importantly, other studies, typically selected for increased survivorship post cold-shock. However, in our study there was very little cold induced mortality. This is further strengthened by the fact that the lifespan of the FSB and FCB populations that were subjected to cold-shock were not different from the longevity of those populations not subjected to cold shock. Thus, it is not surprising that longevity did not evolve in the FSB compared to FCB populations.

In several previous studies, fecundity has responded to selection for cold resistance. Anderson et al. (2005) found that at least two of the three replicates in their selection regime evolved lower fecundity. Watson and Hoffmann (1996) found that cold selected populations had lower fecundity. However, we found no difference in the life-time fecundity of in the FSB relative to the FCB populations. This is in agreement with our earlier, short-term measurement of fecundity in these two populations (Singh et al., 2015). Thus, we found no evidence of a trade-off between evolved cold stress resistance and fecundity.

Increased development time can potentially be a cost in species like *D. melanogaster* that inhabit ephemeral habitats and have to complete their development before the habitat disappears. We did find that the FSB males and females had increased developmental time. However, the magnitude of the increase was very small (∼3-4 hours) and hence we are not sure whether this represented a cost. Increased development time represented an adaptation to increase resource storage that helped in coping stressful conditions. During the late third larval instar stage, *D. melanogaster* larvae feed rapidly and increased their weight exponentially (reviewed in Prasad and Joshi, 2003). An increase of ∼3-4 hours of feeding time during this period drastically increased the amount of resources stored by the larvae. Accordingly, populations of *D. melanogaster* selected for increased starvation and desiccation stress resistance are known to show increased development time and increased body size (Chippindale et al., 1996, 1998). In this study, increased development time represented an adaptation to acquire necessary resources to cope with cold stress.

Body weight at eclosion is often used as a proxy for the amount of resources stored by the larvae. Anderson et al., (2005) and Watson and Hoffmann (1996) found no difference in body size of flies selected for increased cold resistance. In this study, FSB females were heavier at eclosion compared to FCB females. This indicated that FSB females were storing extra/specific nutrients to survive cold-shock. However, there was no difference in body weight between FSB and FCB males. Taken together, this indicated that at least in females, increased development time was likely to be beneficial in aspect of increased resource acquisition. It is also to be noted that in our previous study, females suffered more mortality post cold shock relative to males (Singh et al., 2015).

Absence of any change in lifespan and fecundity of the FSB populations could be because of many reasons. Firstly, the evolved cold-shock resistance ability of the FSB populations might be very cheap. Thus, the resources required to combat the effects of cold stress in our selection regime might be very low. It is a known fact that the flies in our population need to produce active gametes and mate in order to increase egg viability post cold shock. Accordingly, the FSB populations mate more often than the FCB populations post cold-shock (Singh et al., 2015, 2016, Singh and Prasad, 2016). Courtship and mating carry a substantial cost to both males and females (Wedell, 2010). Thus, the costs of evolved cold-shock resistance are expected to be substantial in our selection regime. A second alternative is that the resources are abundant and the FSB populations are able to acquire them as adults. The food used in our selection regime was indeed rich. The larval and adult densities were low. Therefore, it was possible that our flies inhabited resource-rich environment. If this is true, then assays under resource depleted condition should lead to different results. Finally, it is quite possible that the cost of increased cold resistance is paid in a different currency. While we did not find any difference in adult longevity or fecundity, other traits that we have not measured here might have been reduced in the FSB populations. The possible set of such traits include starvation and desiccation resistance.

## CONCLUSIONS

Our findings revealed that there is no life-history trade-offs between increased resistance to cold-shock (in aspect of increased reproductive traits and egg viability post cold shock) with life history traits i.e. the longevity, life time fecundity, larvae to adults survival, and larvae to pre adults developmental time, which indicated that evolved cold stress resistance need not come at a cost of life-history traits. It is quite possible that the cost of increased cold stress resistance is paid in terms of reduced resistance to other stresses.

